# A novel perspective on equal and cross-frequency neural coupling: integration and segregation of the brain networks’ function

**DOI:** 10.1101/2024.05.12.593673

**Authors:** Diego M. Mateos, Jose Luis Perez Velazquez

## Abstract

We introduce a novel perspective in equal and multifrequency coupling derived from considering neuronal synchrony as a possible equivalence relation. The experimental results agree with the theoretical prediction that cross-frequency coupling results in a partition of the brain synchrony state space. We place these results in the framework of the integration and segregation of information in the processing of sensorimotor transformations by the brain cell circuits and propose that equal frequency (1:1) connectivity favours integration of information in the brain whereas cross-frequency coupling (*n:m*) favours segregation. These observations may provide an outlook about how to reconcile the need for stability in the brain’s operations with the requirement for diversity of activity in order to process many sensorimotor transformations simultaneously.

## I. INTRODUCTION

The nervous system functions due to the collective, coordinated activity of cell ensembles, a sort of coordinated activity based on many neurophysiological mechanisms determined by the biophysical, biochemical and anatomical/structural properties of the cells, cell networks and their surrounding environments (e. g. cerebrospinal fluid, epithelial layers). The result of all that bustle is the presence of a variety of rhythms, so-called neural oscillations, observable when neurophysiological activity is recorded using electrodes or sensors that detect field potentials (or their equivalent). These rhythms contain all possible frequencies, and much of the research into how nervous systems ―brains in particular― process information to carry out the wide variety of sensorimotor transformations that constitute the behaviour of an organism relies on the scrutiny of the correlations of those oscillations with behaviours.

The interconnectedness among brain cell circuits (glial cells should not be forgotten, whose activity regulate to some extent that of neurons) results in coherent behaviour emerging at macroscopic (collective) scales arising out of seemingly stochastic activity at small scales [1]. To examine that connectivity ―also termed neural coupling in the literature―, correlations of activity recorded in the brain by various methods are normally used. A common method is to use some sort of synchrony index, and we remark that the strict concepts of synchronization used in physics and mathematics are somewhat relaxed when applied to biological data. Whereas most studies focus on coupling at the same frequency ―which here we will call 1:1, that is, 1:1 frequency locking― examination of cross-frequency coupling (which we will also call *n:m* locking) is becoming popular [2]. A brief technical note: when “coupling” or “connectivity” is mentioned in this text, it is a shorthand for “correlated activity”, as the only coupling/connectivity we can be certain about is that given by the anatomical contacts between cells; the others, so-called functional/effective connectivities, are nothing more than examinations of correlated activity, and there are many texts expounding what is meant by neuronal coupling so we will not delve into it [3]. One reason why assessing cross-frequency coupling is becoming important ―besides the fact that in brain recordings there is almost never a unique frequency rather a mixture of many― is the thought that the interactions among different rhythms provide clues to their function in determining behaviours, some investigators proposing that these different rhythms support distinct components of cognitive acts [4]. Specially, the notion that slower frequencies help integrate faster ones [5] has found empirical support and initiated the view of the communication among neurons via nested rhythms where low frequencies serve as temporal reference for information transfer at higher frequencies [6,7].

Our present study has examined equal and cross-frequency phase synchronization using human electrophysiological recordings and the results suggest a scenario that, according to our knowledge, has never been proposed. Starting from the theoretical consideration of neural synchrony as a possible equivalence relation, we cast our results on equal and multi-frequency locking in the framework of integration and segregation of information in cognition [8, 9]. We propose that equal frequency coupling favours integration from multiple brain regions whereas cross-frequency locking contributes to segregation, or localised sensorimotor transformations in specific brain areas.

The paper is organised as follows. Section II presents the consideration of synchrony as a possible equivalence relation and what this means for neural processing. This is followed by the Methods (section III). Section IV then shows that the prediction from the previous theoretical consideration is found in multiple types of brain recordings. Finally, section V discusses these observations in terms of the known brain dynamics in normal and pathological conditions.

## II. Synchrony as equivalence relation

In neuroscience one talks about connectivity between cells or cell networks. But connectivity means talking about probabilities of connections because this coupling (or connectivity) is determined by both structural (anatomical) and functional (biophysical properties) constraints, and as well by sensory inputs from the environment and the body. Probabilities determine a relationship structure on a system’s dynamical state space, as it is known in the theory of Markov processes (here one talks about transition probabilities), and therefore the related notion of equivalence relation emerges (chapter 3 in [10]). However, while transition probabilities between states that may describe aspects like accessibility from one to another state is an equivalence relation, other measures of correlated neural activity are not necessarily equivalence relations, as explained below. Hence, while connectivity in the anatomical, or structural, sense is an equivalence relation, the functional neural “connectivity” in the neurosciences is not like that of topology ―as mentioned in the introductory paragraphs, functional connectivity is basically assessed as a correlation of activities derived from power, phase or amplitudes of the oscillations, and other observables.

We will discuss in our study the correlations between the phases of the neural oscillations at the mesoscopic/macroscopic level, that is, phase synchronization ―the fundamental importance of phase relations in nervous system function is briefly explained in the Discussion section. We thus choose phase synchrony as an indication of “connectivity”, bearing in mind the aforementioned considerations on neural coupling (and at this point we will drop the quotation marks in the word). Throughout the text we have used phase and frequency locking as equivalent. Although these two, in principle, are not strictly equivalent, in practical terms, in this type of biological analysis phase locking implies to a large degree frequency locking. In fact the equivalence derives from the basic equations for the phase and the frequency of oscillations: phase locking between two oscillations (normally understood as equality of the phase θ) implies θ_1_ = θ_2_, and since the time derivative of the phase is the frequency (ω): dθ_1_/dt = dθ_2_/dt → ω_1_ = ω_2_ then phase locking implies frequency locking. Strictly speaking, in neuroscience the locking condition used for practical purposes is that the difference │θ_1_ - θ_2_│ be relatively constant within a time window —and not the equality of the two phases— which translates to the relative constancy of the difference between the frequencies. Therefore, we use both terms through the text but considering that the common term in this type of studies is ‘cross-frequency (or multi-frequency) coupling’, this expression will appear more often.

Our first question is: is this type of synchrony an equivalence relation? And second: what consequence does this have, if any, for the function of the nervous system? We just briefly mention that an equivalence relation between elements of a set (S) has to satisfy three properties: *reflexivity* (∀ a ∈ S: a ∼ a), *symmetry* (∀ a, b ∈ S: a ∼ b ⇒ b ∼ a) and *transitivity* (∀ a, b, c ∈ S: (a ∼ b) ∧ (b ∼ c) ⇒ a ∼ c), and, if so, this relation establishes clusters of the elements of the set; in other words, the equivalence relation establishes a partition of the state space ―or of the set containing the elements that are being subjected to the relation― into equivalence classes.

The components that are equivalent according to the equivalence relation form an equivalence class ―the different classes are the “cells” of the partition of the set. These classes could consist of only a single point, whereas if there is only one class containing the whole set this is termed the trivial partition, and we will see that this is the case of 1:1 locking. Since in our case we assess phase synchronization between signals, our set is the brain constituted by nerve cell networks (or equivalently, we have a synchrony state space) and we consider whether phase synchrony is an equivalence relation among those elements (neural networks), as it was advanced in other studies [11]. In this perspective, our equivalence classes are the subsets (networks or signals) of the set (the whole montage of signals, or ideally the whole brain) which are equivalent according to our proposed equivalence relation (synchrony); that is, the classes are the signals (recorded from certain brain areas) that are synchronous in either the 1:1 or the *n:m* relation, and as well the signals that are not synchronous to any other (which we call “uncoupled” in this text, these would be the abovementioned single-point equivalence classes). The neurophysiological reason to consider each of the uncoupled neural areas as a subset (in mathematical jargon, a unit set or singleton) is that the signals recorded in these areas originate from neural populations surrounding the electrode or sensor, which may be engaged in certain operations though these might not be connected to other neural networks in different brain regions ―bear in mind that applying strict mathematical notions to biological phenomena requires certain tolerance for flexibility. To clarify: throughout the text we will call “sets” the subsets (equivalence classes in the theoretical perspective) of signals (cortical neural networks because the recording methods we have used detect mostly superficial cortical brain areas) that are synchronous either directly or indirectly via other signals (Figure 1), and as well those that are uncoupled (one-point sets). We note that in these types of analyses one can never know whether the coupling of two networks is due to direct interactions between them or occurs via other networks sending simultaneous inputs to them, but the reasoning detailed below, that will assume direct coupling, remains the same in the case of indirect coupling.

**Figure 1.**
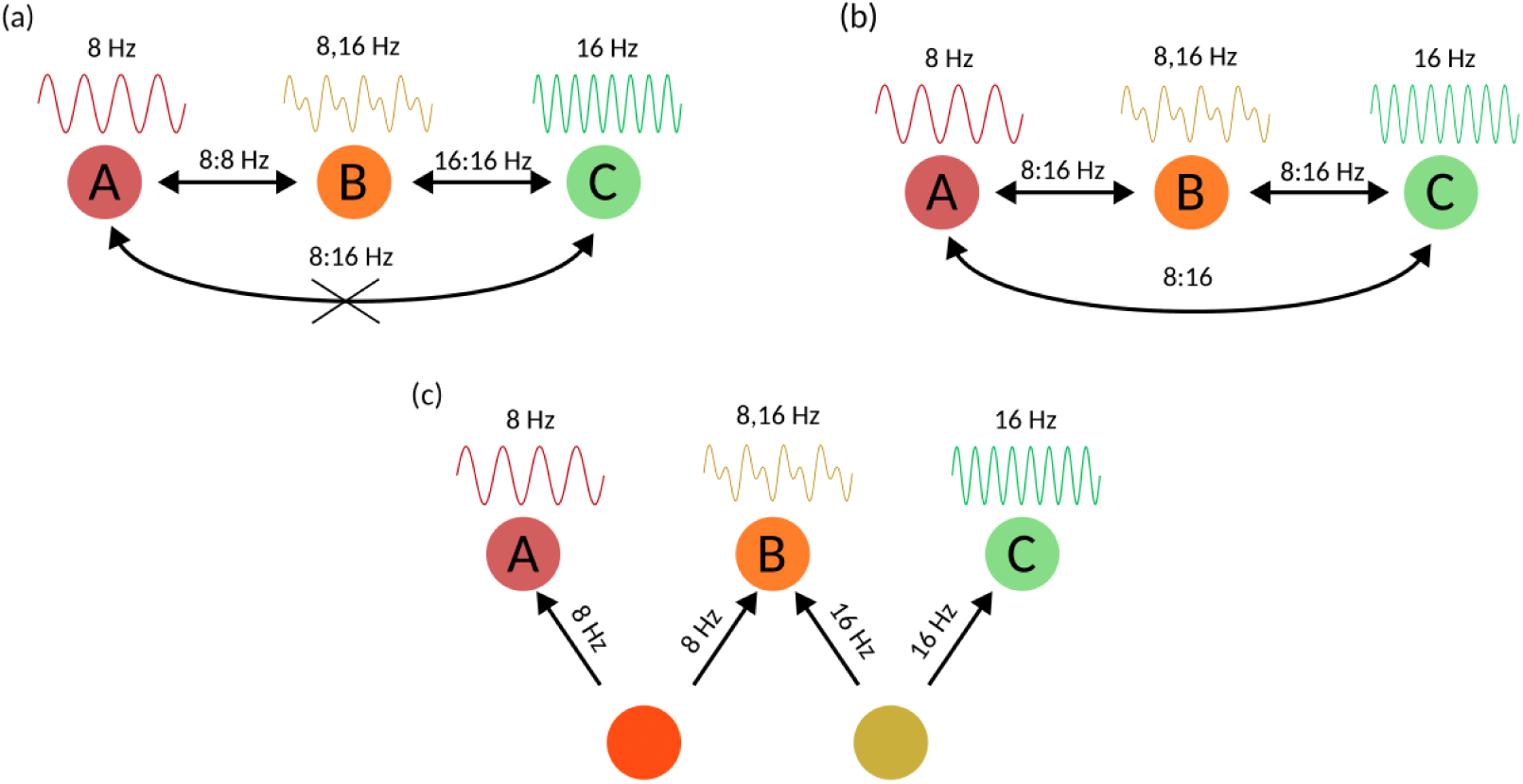
Synchrony as a possible equivalence relation. (**a**) the law of transitivity fails for 1:1 frequency locking; the example shows that at the frequency of 8 Hz there is not a 1:1 relation between neural nets A and C, as explained in the text, hence 1:1 (at a specific frequency) is not an equivalence relation (the waveforms on top of the nets represent the frequencies of activity each net can display, see text for details); (**b**) *n:m* locking for *n*=1 and *m*=2, in this case transitivity is fulfilled, see text for details. (**c**) If the connectivity between the three neural networks is not direct but rather these receive inputs from two other deep nets (from which signals are not recorded), one (left) originating recordings at 8 Hz in the connected superficial nets (A and B) and another (right) producing recordings at 16 Hz (in B and C), same reasoning as in (a) and (b) applies.

As it happens, 1:1 phase locking is not an equivalence relation because while the properties of reflexivity and symmetry are satisfied ―if a cell network A is 1:1 phase locked to another network B, then B is locked to A as well (reflexivity), and of course network A is always synchronous to itself (symmetry)―, transitivity fails. To see this, consider for instance three networks, net A has a pure rhythm of 8 Hz and net B oscillates with two frequencies at 8 and 16 Hz when stimulated by net A, whereas net C oscillates at 16 Hz when it receives input from B, thus A and B can be 1:1 phase locked at 8 Hz, and B and C at 16 Hz, but A and C are not 1:1 synchronous (they would need the same frequency in their rhythms). Figure 1(a) depicts this situation. So at one specific frequency, 1:1 is not transitive; defining this particular frequency to evaluate synchrony is the standard manner that phase synchronization is analysed, and in our studies we call the chosen frequency for each analysis the “central frequency”, as mentioned in Methods.

As to cross-frequency (*n:m*) locking, it is an equivalence relation. It is obviously reflexive, and it satisfies the transitive property because ―continuing with our thought experiment abovementioned― nets A and B will show, say, 1:2 synchrony at central frequency of 8 Hz because B can oscillate at 16 Hz in response to activity at 8 Hz in A, and the same occurs with C in response to receiving signals at 8 Hz from B, hence A and C will be locked at 1:2 too (Figure 2(b)). But symmetry is not that obvious to see, because the phase of one signal at frequency *n* will not be identical to the other phase in the other signal at that same frequency *n*, therefore evaluating phase synchrony *n:m* both ways, say from net A to net B (forward) and vice versa (from B to A, backward), will only be symmetrical if the signals were identical. However, *n:m* does not entail that n<m, it can also be *m:n*, in which case we have the same phases as in the forward case: the phase at frequency *m* in B and phase at frequency *n* in A, thus satisfying symmetry. In fact, one can always find an *n:m* relation between any two frequencies of the many present in the signals —keep in mind in neurophysiological signals we always have a mixture of almost all possible frequencies at which cell networks can display activity—, therefore *n:m* connections are symmetric (in an analogous manner, one can always find frequencies in three networks such that transitivity is satisfied for the *n:m* case). Figure 2(c) shows as well the case of indirect coupling, and one can apply same reasoning as above for each property of the equivalence relation. The consequence of all this is that there is a partition of the “brain set” in the case of cross-frequency connectivity because, as aforementioned, the equivalence relation establishes a partition into equivalence classes of the set containing the elements that are being subjected to the relation; but there is no partition in the case of 1:1 locking since this is not an equivalence relation. The scenario emerging from 1:1 and *n:m* frequency locking with regards to the possible partitioning of the brain cell networks/areas is portrayed in Figure 2.

**Figure 2.**
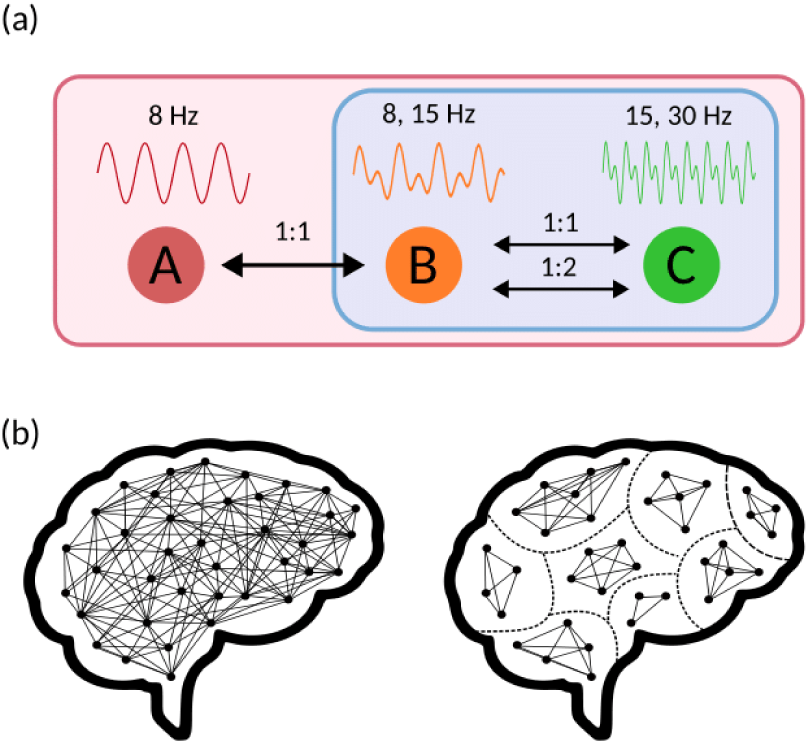
Example of the partition of the neural synchrony state space (**a**) and its graphic representation in the whole brain (**b)**. Three neural nets having activity at 8, 15 and 30 Hz are shown, where information can pass from A to C in a 1:1 locking, A and B oscillating at 8 Hz and B and C at 15 Hz, and thus there is only one set (equivalence class) comprising the three networks (pink rectangle). But for *n:m* locking, only B and C can share information, in this case by 1:2 synchronization (B oscillating at 15 Hz and C at 30 Hz), so we have a partition into two sets (classes), one comprising B and C (blue rectangle) and another containing net A by itself. (**c**) Thus, in 1:1 locking all brain areas are directly or indirectly functionally connected (left) whereas for *n:m* there is a partition (right).

The idea can therefore be advanced that the partition provided by various *n:m* locking ratios could favour sensorimotor transformations in segregated brain areas whereas 1:1 locking could serve for the global integration of that processing made in different sensory or motor areas. Hence, within the framework of integration/segregation of information in cognition [8, 9], 1:1 would facilitate the former and *n:m* the latter.

Now, because of the way in which data analysis is done, described in Methods, *n:m* synchrony is not symmetrical in our analysis, hence the non-symmetrical connectivity matrices (Figure 3). The reason is that, as stated above, we choose one central frequency as the frequency *n* and evaluate only the *n:m* direction (forward and backward) because the neurophysiological question we ask is whether there is transmission of information between two areas, one with activity at *n* Hz and the other responding at *m* Hz. Thus, in the example above of Figure 1 we will find in our analysis 1:2 synchrony from B to C at central frequency of 8 Hz but not from C to B since C does not have that frequency of activity, or even if it had the phase would be different from that of B because the signals are different. If in our analyses we reversed the order and did *m:n* for the case C to B, that is 16 to 8 Hz, this would be a different physiological question: whether in response to 16 Hz activity in C there is 8 Hz activity in B. Thus, we have to continue using *n:m* when evaluating the C to B synchrony in order to be consistent and ask the same neurophysiological question.

**Figure 3.**
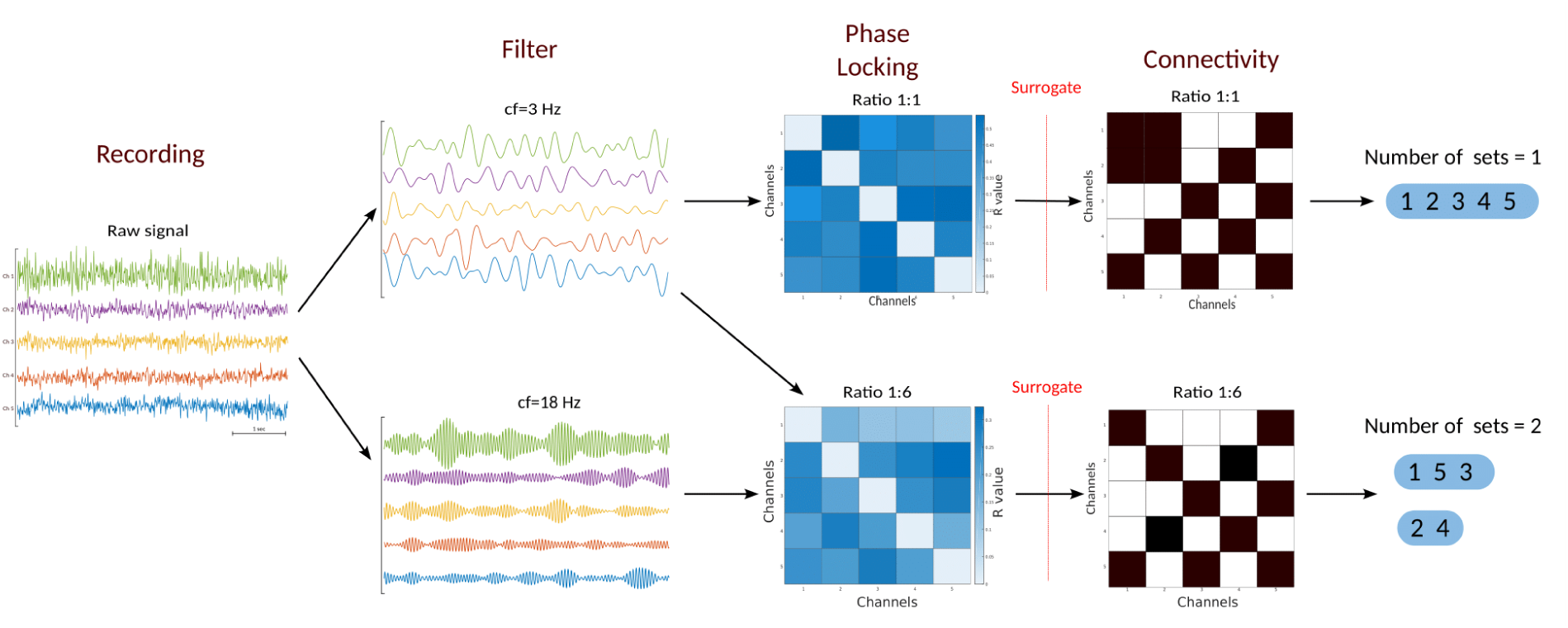
Main analytical procedure common to all analyses. It follows the standard analysis of phase locking, starting with a band-pass filter (+/- 2 Hz) using a central frequency (*cf*), and the final connectivity matrix is Boolean where entries of 1 (black cells in the matrices) indicate that the synchrony index is higher than that of the corresponding entry for the surrogates (see text for details). In the case of cross-frequency analysis (lower row), the connectivity matrix is not symmetric for the reason explained in the text. From these matrices, the number of sets is obtained, the sets representing neural networks that are connected, that is, we assume, as is standard in this type of analysis, that the connected/coupled signals are from neural networks which are involved in a certain sensorimotor transformation (cognitive act). As mentioned in the text, we require that in the non-symmetric matrix both entries for a particular pair of signals to be 1 to define that pair of signals/networks “connected”. As can be inspected in the two sample matrices (not real data) shown, the upper one (1:1) has only one set, and two sets appear in the lower matrix for *n:m* (1:6).

**Figure 4.**
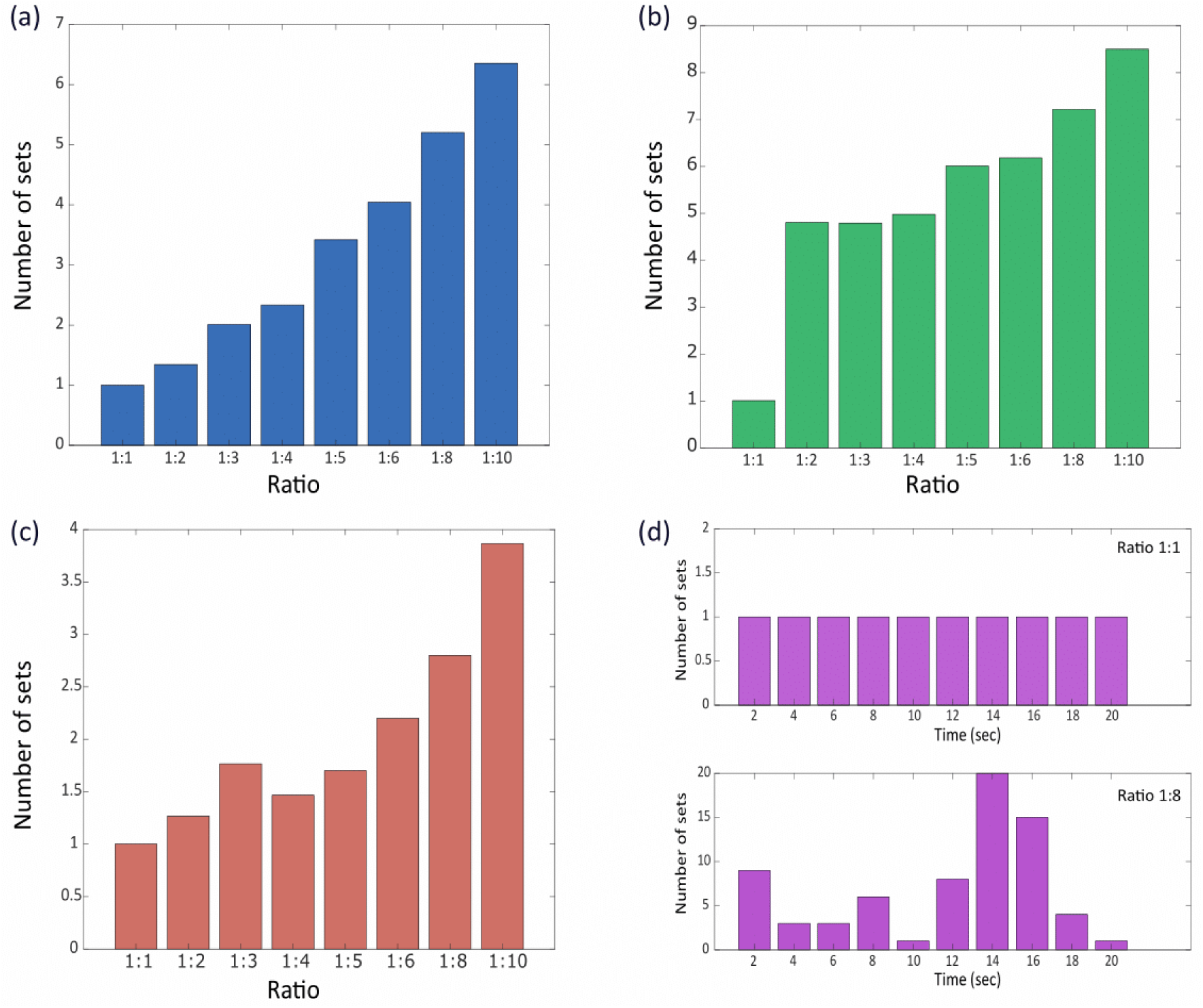
Number of sets found in 1:1 and cross-frequency coupling in **(a)** MEG recordings, **(b)** scalp EEG and **(c)** iEEG signals. Shown are average values for 24 MEG subjects, 111 scalp EEG subjects, and 3 iEEG patients. **(d)** Example, taken from the analysis of one subject, illustrating the variability of the number of sets across ten consecutive time windows (each of 2 seconds) in the case of *n:m* coupling, with the upper figure representing a 1:1 ratio (here there is always 1 set) and the lower figure representing a 1:8 ratio (a central frequency of 4 Hz was used in all four panels).

To understand why the way in which the analysis is done has neurophysiological sense we have to consider the scenario of these three networks where direct reciprocal anatomical (structural) connections exist between A and B and between B and C. It is fair to admit, as we commented above, that we never know for sure the existence of these specific structural connections when performing the analysis, because it may also be the case that the different networks are driven by, say, a fourth network connected to the three of them, so our three nets do not even need to have direct anatomical connectivity to show the functional connectivity that may be directed by the fourth net (Figure 2(c)). These and other considerations arise in the neural synchrony analysis field and are expounded in many papers. For the sake of our argument, let us assume there exist direct reciprocal connections between the three neural nets ―this assumption is reasonable as most of brain areas are reciprocally connected. The neurophysiological question we are trying to answer is whether there is complete transfer of information among the networks, that is, the neural activity proceeds from one net to the next one connected and so on, and *in the reverse direction too*. Therefore we are interested in finding those symmetrical connections otherwise there would be only one direction of information transfer, but to be complete we require it to be in both ways. Again, this requirement is biologically reasonable if we take into consideration that almost all anatomical connections in the nervous system are recurrent ―everything is connected directly or indirectly to everything in the brain. This organization has received other names such as re-entry, a process whereby brain regions concurrently stimulate and are stimulated by each other supporting the synchronization of neural firing. Therefore, when we choose, say, 8 Hz as central frequency —continuing with the above example— we are interested to ensure that whatever information is transferred at 8 Hz proceeds both ways. Later we can try another central frequency like 16 Hz, and so on, to explore the arrangement/configurations of connections at that frequency in the various *n:m* relations, including 1:1. So, to insist again: if we were to analyse the 1:2 transfer of activity from B to C as 8:16 Hz (forward), and then at 16:8 Hz (that is, 2:1) when going from C to B (backward), we would be asking two different neurophysiological questions.

## III. METHODS

### A. Electrophysiological recordings

Brain recordings were analysed from a total of 137 adult subjects. Two open databases of healthy participants included 20 magnetoencephalographic (MEG) recordings from the Human Connectome Database (db.humanconnectome.org) and a group of 109 scalp electroencephalographic (EEG) recordings from the EEG Motor Movement/Imagery Dataset [12], the former recorded at a sampling frequency of 508 Hz and the latter at 160 Hz (with 64 electrodes using the international 10-10 system). In addition, other 8 recordings that we used in the past were analysed in the present study: three epileptic patients recorded with MEG, three epilepsy patients studied with simultaneous intracranial EEG (iEEG) and scalp EEG, and two nonepileptic subjects studied with scalp EEG, all these subjects have been described in previous publications [13, 14]. The number of sensors in each MEG or EEG montage varied from patient to patient and it is specified, when needed, in the text, and as well the sampling frequency varied from 200 to 625 Hz, differences that were taken into consideration for the data analyses.

### B. Data analyses

The recordings corresponding to the two external datasets were initially processed with a bandpass filter in the frequency range of 0.5–150 Hz, and a notch filter at 50 Hz was applied to remove power line interference. In order to avoid the potential effects on synchronization of the common reference electrode used in scalp EEG and of volume conduction, the scalp EEG recordings were pre-processed with the current source density (CSD) toolbox (MATLAB implementation [15]) to compute the scalp surface Laplacian or current source density estimates for surface potentials using a spherical spline algorithm [16]. To further diminish signal summation and volume conduction, one half of the sensors of the high-density scalp EEG and MEG montages were used in the analysis, the retained electrodes being equally distributed on the scalp. A variable length of the recordings were used for the different analyses, ranging from 10 to 60 seconds, and these segments were divided into consecutive windows ranging from 2 to 30 seconds; and when needed, longer segments of 5 minutes were examined (all these details specified in the text as needed).

Our analytical procedures start with the calculation of phase synchronization between two signals (described in detail in [13]). Thus, a phase synchrony index was calculated for all possible pairwise signal combinations, for which we use the standard procedure of estimating phase differences between two signals from the instantaneous phases extracted using the analytic signal concept via the Hilbert transform. To compute the synchrony index, several central frequencies, as specified in the text and figure legends, were chosen with a bandpass filter of ±2 Hz; hence, for one particular value of the central frequency *cf*, we end up with a signal with frequencies c*f*±2 Hz. This filter is necessary because to have a clear physical meaning of the instantaneous phase the signal has to be a narrow-band signal (for details on the analytic signal approach see appendix A2 in [17]). The *phase synchrony index* R between two signals is calculated from the average difference between the phases of the oscillations using a 1-second running window (this window to compute R should not be confused with the window of the signals we use in the rest of the analyses) using the mean phase synchrony statistic which is a measure of phase locking and is defined as 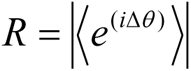 where Δθ is the phase difference between two signals, also termed relative phase [18]. The more stable the Δθ in the specific time window (1 second), the closer the R value is to 1 (R values are between 0 and 1). The following analytical procedures have been described in [19] and in [20]. Briefly, the synchrony indices obtained in this manner for all pairwise combinations of signals in a specific electrophysiological montage gave us a matrix whose entries are the average values of the index R for each pairwise configuration during the time period investigated. From this matrix, a Boolean (binary) connectivity matrix is calculated, with entry of 1 if the corresponding synchrony index is higher than a threshold, and 0 if lower. The threshold is obtained from the mean R given by (twenty) phase randomised surrogates of the original signals. For this purpose, another phase synchrony matrix of the surrogates was obtained applying the aforementioned method, and each entry in this surrogate matrix was compared with the corresponding entry in the original matrix of the signals, resulting in the binary “connectivity” matrix where a value of 1 is assigned to an entry if the R value of the signals is larger than that of the corresponding surrogates. Please note that we do not use an average of all pairwise surrogate indices to define the threshold, although we note that the results were qualitatively similar if a grand average threshold was used, but we think it is more accurate to use the threshold derived from the surrogates for each specific pair of signals. The reason is that although the surrogates are in principle randomised time series, in reality they carry some features of the original signals, therefore we observe that the synchrony index between surrogates of neighbouring signals is higher than that between surrogates of distanced signals, probably because the known phenomenon of signal summation in adjacent sensors; hence, comparing each pairwise combination of signals with their corresponding surrogates is probably better than using a grand average among all surrogate pairs, although we insist the results were qualitatively analogous using the latter.

The condition for cross-frequency, or *n:m* phase locking is analogous to the previous case for 1:1 locking, the relative phase is now Δθ*_n,m_*(t) = *n*θ_1_(t) - *m*θ_2_(t), *n* and *m* are integers, θ_1,2_ are the phases of the two oscillators [21]. Our method to compute *n:m* synchrony and the Boolean connectivity matrix then proceeds identically as described above for the 1:1 case by extraction of the instantaneous phases of the signals at the required frequency ratios; in our studies we used *n*=1 and *m* ranged from 1 to 10, the particular *n:m* ratios studied are specified in the Results section as needed. The only disparity with the 1:1 case is that while the synchrony index matrix is symmetrical for 1:1 it is not so for *n:m* due to the different R values when the phase synchrony is computed in one direction (forward, as described in section II) and the reverse (backward), and this occurs because the instantaneous phase at frequency *n,* or *m*, is different for signal 1 and signal 2 and, obviously, the phase difference will only be identical if n=m. Because we consider two signals “connected” if they are so in both directions for the reasons detailed in section II, then in the case of the *n:m* matrix we have to evaluate both entries for each pair of signals in the corresponding binary matrix. Once the binary connectivity matrices were obtained, sets of connected signals were found based on the concept introduced in section II, which in simple words states that "if signal 1 is connected to signal 2, and this one is connected to signal 3, then 1 is also connected to 3". Figure 3 summarises all these analytical procedures.

The number of configurations of signal connections was estimated as described in [19, 20]. Briefly, the number of configurations of the connections found in the Boolean connectivity matrices is 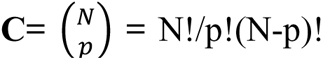 where **C** is the number of possible combinations of the ***p*** pairs found connected, and *N* is the total number of possible pairs of signals given a specific channel montage, that is, 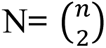 for *n* sensors/signals in the montage. In sum, **C** represents in how many ways our experimental observation of *p* connected pairs can be arranged given a maximum of *N* possible pairs. Due to the large numbers of connected signals and possible pairs, the binomial coefficient cannot be computed in most cases, so we take the logarithm and use the Stirling approximation to estimate the number **C** of connectivity configurations, that after some basic algebra becomes ln(C) = N × ln(N/(N − p)) – p × ln(p/ (N − p)). Finally, to visualise the result graphically, the values of ln(C) are plotted against N which results in an inverted U, and the experimental points corresponding to the ln(C) for each *p* obtained are graphed on that curve (see Figure 5 and those in [19, 20]).

**Figure 5.**
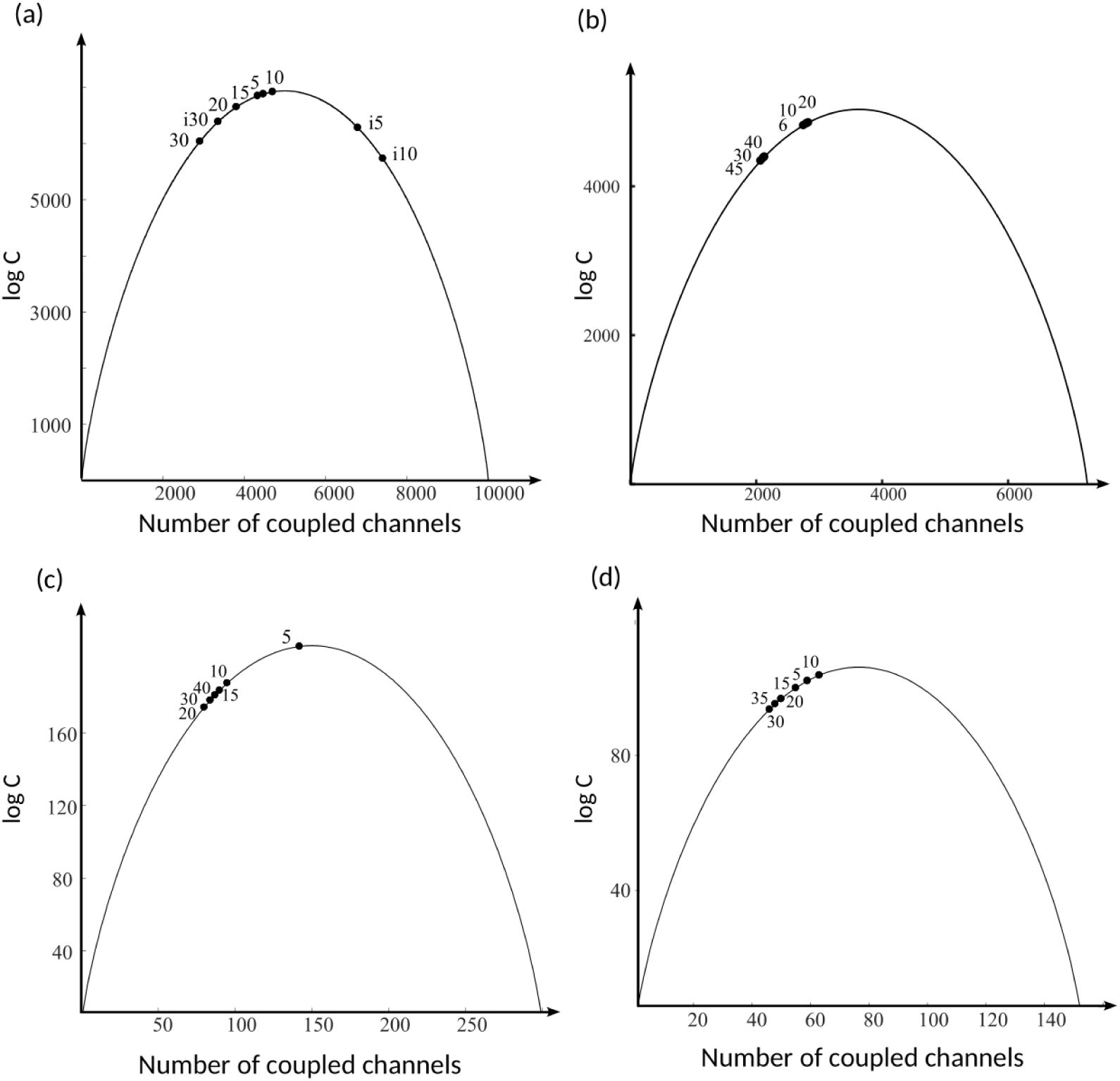
The logarithm of the number of configurations of pairwise connected signals at different frequencies. The continuous curves are obtained from the log(C) equation shown in Methods and represent the values of log(C) for all possible numbers of pairwise combinations of signals, while the dots represent the values for the different central frequencies written next to the dots. Shown are the results in a MEG recording of a patient with generalised seizures **(a)**, another MEG in a healthy subject **(b)**, a scalp EEG recording **(c)**, and a iEEG recording **(d)**. In all cases, lower frequencies tend to be closer to the top of the curve, where the maximum number of configurations is found. The farther from the top, the fewer number of connectivity configurations. In **(a)**, the data labelled i5, i10 and i30 represent the results from the ictal event (generalised epileptic seizure that resulted in an unresponsive state) in this patient at those three frequencies, and, as reported previously [19, 20], the numbers of connectivity configurations are lower than those corresponding to the same frequencies during a normal state (without seizures) in that patient, except for 30 Hz.

## IV. RESULTS

Based on the reasoning of section II, we have examined the partition of the neural synchrony state space in 1:1 and cross-frequency locking using the three types of neurophysiological recordings stated in Methods: 23 MEGs, 3 iEEGs and 114 scalp EEGs. Evaluating signals in three different techniques ensures the results are robust and do not depend on the methodology of the data acquisition, because each method has its drawbacks. All recordings were taken during the so-called resting condition ―that is, when subjects are not performing any specific behavioural task― except for the two patients in coma; in addition to these two patients, other four with generalised epileptic seizures (three recorded with MEG and one with simultaneous scalp/iEEG) gave us the opportunity to examine possible differences in terms of the partition of the neural network space during unconscious states as the generalised ictal events were accompanied by unresponsiveness, so we have the two patients in coma and the four with seizures as samples of unconscious states. To compare these pathological unconscious states with the normal unconscious state of slow wave sleep, we analysed four individuals in deep slow wave sleep (stages 3-4), two recorded with iEEG and two with scalp EEG.

The same results were found in all conditions and recording methodologies: there is partition in the case of cross-frequency coupling and no partition for 1:1 coupling. The number of sets in the partition for *n:m* locking varied, but it was always one for 1:1 locking, and the composition of each set in the *n:m* case was variable from window to window (signals were segmented into non-overlapping 2-sec windows, as described Methods). Figure 4 shows representative results that were qualitatively identical in the 3 types of recordings evaluated in this study. We used 1 and 2-second windows (the majority were done using 2-sec) because we tried to be close to a reasonable cognitive temporal scale when sensorimotor transformations among neural circuits take place, which normally ranges between 50 and 500 ms; windows shorter than 1 or 2 seconds would not be adequate due to the constraints of the phase synchrony analytical procedures. We note that very long windows, from 20 seconds to 5 minutes, resulted in no partition at any *n:m* examined, due to the high probability that in this long length all signals will be directly or indirectly connected: the longer the temporal scale the higher the probability that the average synchrony index for a particular pair of signals will be higher than that of their surrogate pair because the index for the surrogates decreases with longer timescales due to their random nature; and, at the same time, the synchrony between the original signals increases with the length of the time window considered, which is to be expected in brain signals because the coordinated activity of neural circuits favours the tendency towards a more correlated activity if longer time windows are considered, and note that this is the opposite of the surrogates’ tendency as mentioned above because the surrogates are random time series that supposedly present little or no coordinated (correlated) activity. However, we would like to note that the increase in the synchrony index with the length of the signal reaches a plateau, a value that remains more or less constant, after a certain length (it stands to reason it cannot continuously grow until reaching its maximum value of 1). In our analyses, this time length was about 15 -20 s. We comment on this because this may represent the physiological tendency towards maintaining a steady state. This tendency can be seen at any level, for example at the molecular level the distribution of Na^+^ and K^+^ ions inside and outside of cells remains relatively constant in the long run, although it is briefly altered during an action potential; hence, looking at short time scales we will see instances with very different ion distributions but the average distribution will be nearly constant inspecting longer scales. In previous studies, the different views on the short and long time scales in neural connectivity were explored, resulting in high variability at shorter time scales and a relatively steady state at longer ones [20].

While the two pathological cases of epileptic seizures and coma here investigated presented same results as those of the healthy population, there are some crucial differences in another aspect of the connectivity; this feature is the number of configurations of connections, namely, there are more configurations in the healthy recordings than during unconscious states of seizures and coma as reported in [19, 20] (see also Figure 5(a) where the results obtained in one seizure are shown). This occurs because most of times there is enhanced synchrony ―more signals connected― in these pathological situations which lowers the number of configurations of the pairwise connected signals (the number of configurations of signal connections was calculated as explained in Methods). As well, a main difference in the coma recordings as compared with the non-pathological ones was that most signals were uncoupled in *n:m* locking, perhaps reflecting the uncoupling observed in unconscious states caused by the brain damage in coma, vegetative or minimally conscious states [22, 23, 24, 25].

The presence of one set incorporating all signals, in the case of 1:1 locking, occurred at all frequencies studied spanning from 3 to 45 Hz. We have also seen that the same occurs in pathological cases during epileptic seizures or coma where consciousness is absent or greatly reduced, but we know that the difference between the whole connected set of normal consciousness and that of the pathological cases is that there is a greater number of connectivity configurations in the former [19, 20], hence the possibility arises that the distinct frequencies could present different number of connectivity configurations in spite of all having a whole connected set of brain areas in 1:1 locking. To test this, we computed the (approximate) number of connectivity configurations using the Stirling approximation described in Methods. Figure 5 depicts some examples (four subjects with different recording methods) illustrating the main result: larger number of configurations of network connections can be seen at lower frequencies. The dots in the figure are the number of configurations taken from the average of connected pairs in signal segments ranging from 10 to 30 seconds that were divided into 2-sec consecutive windows as in the frequency locking analysis presented above. We note again that, as occurred in the previous analysis, there were no apparent differences at any frequencies analysed using windows of 10 sec or longer, so this is, again, an indication that this phenomenon is associated with reasonable cognitive timescales (that is, short), rather than with global and temporally extended behavioural states. The difference in the number of configurations between low and higher frequencies is clearer at frequencies over 15-20 Hz: whereas at low frequencies (3 to 10 Hz), the number of configurations was similar, the decrease in this number was becoming prominent at frequencies over 15 Hz in most subjects analysed. Specifically, this phenomenon was observed in 6 of 6 MEGs, in 4 of 4 iEEGs, and in 5 of 7 scalp EEGs. The scalp EEGs that did not present this phenomenon were characterised by high power at frequencies between 20 and 40 Hz, possibly due to unusually high scalp muscle activity which was confirmed by the neurophysiologist’s report. We note that it is known that neither the CSD procedure nor any other (e.g. inverse source reconstruction) can eliminate completely scalp muscle activity [26, 27], and even other common problems like volume conduction cannot be eliminated by source reconstruction procedures [28]. This observation of larger number of connectivity configurations for lower frequencies goes along the lines of the importance of these slow frequencies in brain activity and is discussed in the next section.

To finalise this section, we present the results examining the possible partition (or its absence) of the synchrony state space using artificial signals. We credit one of our referees for suggesting this analysis. One may cast doubt that our results represent a specific feature of neurobiological signals, suspecting that it is a generic property of any signal regardless of origin. This doubt is legitimate but only in part. It is clear that depending on how correlated the signals in a system are —signals of any kind, not necessarily neurobiological— the synchrony state space will be partitioned or not. We thus created an “arbitrary” system using surrogates of the original signals, that is, random time series, but maintaining same number of signals (channels) and same sampling frequencies as the originals. We subjected these artificial signals to an identical analysis done with the originals as abovementioned. The results were similar to those of the original signals in that for *n:m* locking there was a partition of the synchronization space — although in this case the numbers of sets were higher— but, unlike with the original signals, in the 1:1 locking condition sometimes there was a partition, which we never saw with the original systems. These results are expected because they represent the low correlations in the random, artificial signals, thus more sets appear for multifrequency coupling and sometimes there is not enough correlation to maintain all connected in the 1:1 case. If we had increased the correlation among our artificial signals, then they would have looked more closely like those of neurobiological origin —after all we have to remember that neurophysiological signals at the meso/macroscale represent noisy, quasi-random time series with a very large number of frequencies all mixed together— and therefore the results would have been closer to those we obtained with the original system. We therefore think that it is a generic property of any system that, depending on the nature of its signals (specifically their correlations), the synchrony state space (if one such space can be defined by that system) will show a partition for *n:m* locking (with n ≠ m). The important matter is that *this result has to be interpreted for each system*. In our case —brain research— we have framed it within the integration/segregation notion of brain information processing. The interpretation for other systems could be of interest. For example, one can talk about the association of multifrequency relations in the activity of heart muscle with cardiac arrhythmias [29, 30, 31], where it has been shown that multifrequency oscillations create a unstable prefibrillatory condition; we could speculate that the partition of the, in this case, ‘cardiac muscle state space’ due to those multifrequency relations favours the instability of the required muscle coordinated activity for proper heart function.

## IV. DISCUSSION

Our purpose in this study was not to evaluate the fine structure of cross-frequency coupling in terms of specific anatomical regions or preferred frequency locking ratios but rather to inspect one basic, general feature derived from the theoretical consideration of synchrony as a possible equivalence relation. This global dynamic feature consists in partitioning the brain into multiple sets of connected cell networks in the case of multi-frequency coupling, whereas there is one set encompassing all brain areas that display 1:1 locking. The fact that the results were qualitatively the same in the three recording methods evaluated suggests that these observations do not represent an artefact of the recording technique. We place these results in the framework of the integration and segregation of information in the processing of sensorimotor transformations by the brain cell circuits [8, 9] and propose the novel viewpoint in the field of cross-frequency connectivity that equal frequency locking (1:1) favours integration of information processed from multiple brain regions whereas cross-frequency locking (*n:m*) contributes to segregation, or localised processing in specific brain areas. Each set, or equivalence class, groups the cell nets that share a common frequency (which is needed to be frequency locked), and since these frequencies evolve over time during brain activity, the networks can change from one equivalence class to another ―the sets of coupled brain areas change continuously as we found in our study. This reflects the fluctuations of brain activity that are fundamental for a healthy brain function, as decreased variability in neural activity and synchrony in particular has been associated with pathological cases [32, 33].

Our results are in line with theoretical/computational studies on brain cortical areas interacting through phase modulations; particularly, “…an entire cortex can be dynamically partitioned into interwoven but relatively independent ensembles […] A cortical oscillator may participate in different ensembles by changing its frequency” [34]; this is precisely what was found in the case of *n:m* coupling. Other reports on “chimera” states where there is a breakup of synchronization of a population of oscillators into diverse subpopulations [35] find empirical support in our observations. The results indicating that synchronous networks are biased towards internal information processing but incoherent ones are biased towards the processing of external information [36] could be interpreted using our observations of the equal frequency connectivity ―without partitioning the synchrony space― as being the “internal” mode and the cross-frequency coupling ―with the consequent partitioning― as the external information processing mode because each cortical region tends to be specialised to process one sensory or motor aspect of the behaviour. It should be remarked too that from an anatomical/structural level, studies on brain network parcellation using neuroimaging data have indicated the rich structure of these parcellations [37, 38].

Since in the case of 1:1 locking all brain areas are directly or indirectly connected, one could hypothesise that, in principle (although this cannot be known with the current neuroscientific technologies), any information can travel from one brain area to any other, thus favouring integration of information. On the other hand, multifrequency locking, by imposing a partition of the brain synchrony space, promotes the specific processing of sensory inputs and motor programmes in particular brain areas devoted to these tasks ―e.g., the auditory cortex processing auditory inputs or area M1 (primary motor cortex) processing motor actions. This partition also allows the processing of multiple tasks simultaneously. And because all these dynamic modes operate at the same time in a non-pathological brain, the simultaneous integration and segregation of information is guaranteed.

Other studies have assessed the fine structure of multiple locking in terms of the anatomical areas that display cross-frequency connections. As a representative example, one study has shown that large-scale networks of cross-frequency connectivity is characteristic of resting state brain activity, and the observed coupling of slow and fast oscillations between anterior and posterior parts of the brain indicated that this type of coupling coordinates intrinsic and extrinsic processing modes [39], along the ideas in [36]. In fact, there is an extensive literature on cross-frequency connectivity (reviewed in [40]) due to the increasing interest in addressing this most fundamental aspect of brain recordings: that a unique frequency in the signals is never observed, even in very pathological cases like coma what is recorded is a mixture of frequencies. Therefore, investigating the interactions among these multiple rhythms may provide clues to their function in determining behaviours [4]. One popular idea is that slower frequencies help integrate faster ones [5, 41], thus low frequencies have been proposed to serve as a temporal reference for information transfer at higher frequencies [6, 7]. Of note, even a practical use of cross-frequency coupling has been proposed using noninvasive neurostimulation to improve working memory [42]. Although we found similar global features occurring in the healthy recordings and in the two pathologies studied here, epileptic seizures and coma, other studies have scrutinised in more detail cross-frequency coupling in diseased (psychiatric) and normal states in terms of anatomical distribution and frequencies involved, reviewed in [43].

Importantly, a fundamental feature differentiating the healthy and the pathological brain dynamics may not be the partition (or lack) of the synchrony space but rather the number of connectivity configurations: maximising this number is associated with conscious as opposed to unconscious states [44]. Interestingly, we see a similar tendency of maximising connectivity configurations for low frequencies, in spite of finding a whole connected set of signals in 1:1 locking for any frequency that we studied, from 3 to 45 Hz. This occurs because at low frequencies there is, normally, enhanced synchronization, which results in more connected pairs of signals and therefore the number of configurations move closer to the top of the inverted U; we can see this clearly in Fig. 5, notice at higher frequencies the data points are more displaced to the left of the curve because of a lower number of connected pairs (the x-axis in the graphs), but due to higher synchrony the lower frequencies display a larger number of connected pairs and thus more connectivity configurations. But too much synchrony, like that during seizures, results in too many connected pairs and therefore lower number of connectivity configurations, as we see clearly in Fig. 5(a) where two of the three ictal frequencies analysed (5 and 10 Hz) fall to the right of the top of curve. These observations are along the lines of the proposal that a widespread and long-lasting neural synchrony is normally associated with diseases [45].

We have used phase synchronization because of the fundamental role of phase relations in the activity of the nervous system [46, 47]. Phase relations provide an effective means to control the timing of neuronal firing [48]. This is evident just by considering what the neurophysiological signals represent: the amplitudes of the oscillations reflect the synaptic and action potentials in neurons ―although due to the very short duration of action potentials it is mainly the synaptic activity which is recorded in EEGs. Those synaptic inputs may arrive either in synchrony or stochastically, the former (to be more precise, especially in-phase synchrony) enhancing the probability that the receiving neurons will cross the threshold for action potential generation and therefore will fire spikes and spread the activity to other connected neural nets. Perhaps the clearest examples of how the subtle changes in neuronal synchrony patterns determine the final outcome of specific behaviours are found in the case of the motor system where the evidence for cooperation among neurons via synchronization of their activities is manifest (reviewed in [49]). Nevertheless, other analytical methods have been used to characterise cross-frequency connectivity, e.g. phase-amplitude coupling [50]. Considering that classical synchronisation theory distinguishes between mutual synchronisation of self-sustained periodic oscillators and forced synchronisation by driving forces, in the case of the mammalian nervous system it seems more plausible that the latter is the main phenomenon because one neuron receives a multitude of synaptic inputs from many cells, and thus the input it receives is strong, although mutual synchronisation has been described too [51].

In general, our results may provide a perspective about how to reconcile the need for stability in the brain’s operations with the requirement for diversity of activity in order to process many sensorimotor transformations almost simultaneously. The global pattern of 1:1 locking conceivably imposes some stability in terms of integration of a wide variety of sensorimotor processing in many brain regions, although the connectivity configurations are high enough so that a wide variety of connectivity patterns —especially mediated by lower frequencies which modulate the local synchronization at higher frequencies [5, 6, 7, 41]— emerge from moment to moment to help the organism navigate its world. This idea is supported by studies like those examining the functional connectivity of large scale neural activity that found transient coactivations of localised cortical regions at different times during the global propagation of the waves across all cortex [52], which suggests that the spatial distribution of the localised activations is embedded in the phase of the global waves; as well, the common proposal that cross-frequency connectivity facilitates flexible coordination of the spatio-temporal neuronal activity [40] is supported by our observations.

Our study has the typical limitations of empirical studies based on theoretical considerations: it represents an attempt to translate (strict) theoretical concepts ―equivalence relations― to biological activity and all this constrained by the analytical methods, therefore several assumptions must be made. Our main assumptions have been described in section III as these relate to the conceptualisation of neural synchrony as an equivalence relation [11] considering the physiological facts about neuronal activity and the analytical methods to assess synchrony; still, these are reasonable assumptions which led us to find a main difference between the global dynamics of equal and cross-frequency coupling. We have as well remarked that the fragmentation of the synchronization state space is a generic property of any system with signals that can be studied using some sort of correlated activity, and the interpretation of such partitions (or their absence) must be evaluated in each specific system being studied. Our proposal for this novel perspective on coupling among the neurophysiological rhythms may be useful to continue characterising how neural activity unfolds continuously to generate the variety of behaviours and may have possible approaches to understand pathological neural activity.

